# Dynamic metabolic modeling of ATP allocation during viral infection

**DOI:** 10.1101/2024.11.12.623198

**Authors:** Alvin Lu, Ilija Dukovski, Daniel Segrè

**Affiliations:** Bioinformatics Program, Faculty of Computing and Data Sciences, Boston University, Boston, MA, USA; Yale University, New Haven, CT, USA; Biological Design Center, Boston University, Boston, MA, USA; Center for Advanced Interdisciplinary Research, Ss. Cyril and Methodius University, Skopje, N. Macedonia; Department of Physics, Boston University, Boston, MA, USA; Department of Biomedical Engineering, Boston University, Boston, MA, USA; Department of Biology, Boston University, Boston, MA, USA

## Abstract

Viral pathogens, like SARS-CoV-2, hijack the host’s macromolecular production machinery, imposing an energetic burden that is distributed across cellular metabolism. To explore the dynamic metabolic tension between the host’s survival and viral replication, we developed a computational framework that uses genome-scale models to perform dynamic Flux Balance Analysis of human cell metabolism during virus infections. Relative to previous models, our framework addresses the physiology of viral infections of non-proliferating host cells through two new features. First, by incorporating the lipid content of SARS-CoV-2 biomass, we discovered activation of previously overlooked pathways giving rise to new predictions of possible drug targets. Furthermore, we introduce a dynamic model that simulates the partitioning of resources between the virus and the host cell, capturing the extent to which the competition depletes the human cells from essential ATP. By incorporating viral dynamics into our COMETS framework for spatio-temporal modeling of metabolism, we provide a mechanistic, dynamic and generalizable starting point for bridging systems biology modeling with viral pathogenesis. This framework could be extended to broadly incorporate phage dynamics in microbial systems and ecosystems.

## Introduction

The recent COVID-19 pandemic has demonstrated the need for fast and efficient methods of encoding complex biological information, forecasting the outcome of different virus variants, and implementing scalable models that can help prioritize the screening of possible drugs and drug targets. While there is currently no “*in silico*” substitute for the long and expensive processes needed for identifying, developing and testing vaccines and therapies, there is increasing interest in the exploration of computational approaches that can help accelerate specific aspects of this process and prioritize avenues for drug discovery.^1–4^

Viral infections of host cells constitute complex multiscale biological processes, whose dynamics cannot be easily grasped through simple intuition, but it is rather best explored with the help of mathematical models. These models may cover a broad range of spatial and temporal scales, from the dynamics of individual proteins and reactions,^1,5–8^ to population-level epidemiological models.^9^ Ideally, a mathematical model would facilitate cross-scale inferences, e.g. by predicting the fate of an infected cell based on the collective behavior of the thousands of reactions that cells and viruses struggle to control. One such approach, genome-scale stoichiometric modeling, is specifically focused on understanding the allocation of metabolic resources in a cell, and has been proposed as a tool to help decipher the physiological changes associated with viral infections.^1,5–8^ This approach involves two fundamental steps: the first (metabolic reconstruction) is the encoding of all metabolic reactions of an organism in a formal mathematical representation; the second (phenotype prediction) is the usage of this mathematical representation, coupled with environmental information, to provide mechanistic predictions of the behavior of the organism.

A frequently used method for this second step is Flux Balance Analysis (FBA), which uses efficient linear programming algorithms to solve the problem of resource allocation in the cell at steady state, by assuming that the fluxes have evolved towards a predictable optimal state.^10–14^ An objective function frequently used for these calculations is biomass production, equivalent to the assumption that in managing its complex balance of supply (nutrients) and demand (biomass components), an organism will achieve a state of maximal growth capacity.^10–14^ Genome scale metabolic (GEM) models were initially developed and tested in bacterial metabolism,^11–14^ and have become increasingly useful for studying microbes and microbial communities, as well as human metabolism.^11–14^ Human cell reconstructions with increasing degrees of completeness and accuracy have been gradually introduced, including Recon1,^15^ RECON3D,^16^ and the Human Metabolic Reaction (HMR) database.^17^ These human models have been used for a multitude of studies, including, recently, analyses of how cell-virus metabolic interactions affect host metabolism under viral infection.^1–5^ Upon infection, a virus typically modulates the metabolic activity of its host cell and reallocates resources for its own proliferation, competing against host processes (**Fig. 1**). This competition for resources makes FBA a potentially valuable way of investigating the tug of war between the host and virus under given supply conditions.

**Figure 1:**
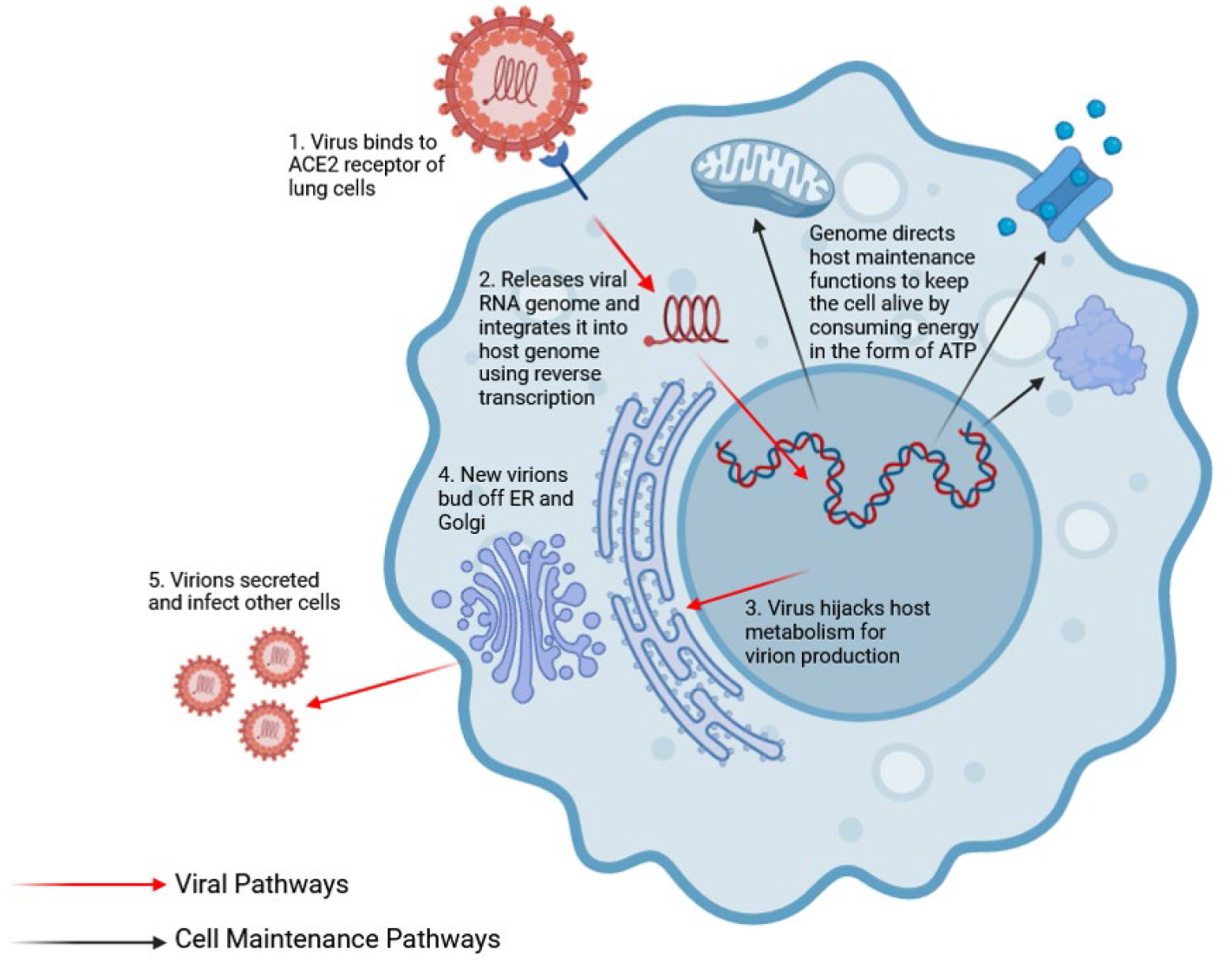
Viruses rely on host metabolism to reproduce by hijacking cellular processes and reappropriating resources for its own replication. The schematic shows the process of SARS-CoV-2 infection. Red arrows indicate viral pathways of replication, from when the virion binds to the cell membrane and releases its RNA to the synthesis of new virions in the ER and Golgi. Meanwhile, the black arrows indicate cell maintenance pathways, which consumes energy to keep the host alive (such as synthesizing its own proteins, cellular transport, etc). This creates competition between the host and the virus.

Most papers that utilize GEMs for modeling virus infections have followed a framework that consists of two major steps, akin to the basic steps employed in FBA: the first step consists in modifying the genome scale reconstruction itself by adding a supplementary biomass-producing reaction, which encodes viral reproduction.^1–3,5^ The second step is a procedure to simulate the interplay between host and viral metabolism. A frequent approach for this second step has been to maximize a linear combination of the reactions representing growth of the host and virus biomass, trying to capture the concurrent activity of the two processes, and the competition between host and virus.^1–3,5^ While valuable, this approach has a number of limitations which we address in the current work. First, these existing approaches lack a temporal component; to the best of our knowledge, all metabolic genome-scale models for host-virus interactions developed so far only employ standard FBA, which computes reaction rates at a single moment and does not track cell growth over time. Thus, they do not address factors such as the dynamics of nutrients’ depletion or the effect of the host dying off over time due to virus proliferation. Second, most human cells infected by viruses such as SARS-CoV-2 (e.g. alveolar macrophages and lung epithelial cells) are not actively growing, and optimizing their biomass growth rate may not be accurately capturing their natural state. Most healthy cells are only maintaining their metabolic state through energy consumption, primarily in the form of ATP hydrolysis. The infected cells, on the other hand, are producing virus biomass, a process that also requires ATP consumption. If the virus’ demand for ATP brings the levels below the amount needed by host cells to survive (maintenance flux), the expectation is that these cells would not be viable any more. Third, employing a linear combination of objectives would generally converge to either the virus or the human cell “winning” all of the resources, with no middle ground where human tissue may struggle but survive, while the virus simultaneously replicates at a lower rate. Notably, recent experimental work on viral dynamics has specifically highlighted the importance of energy currency competition for understanding takeover of host cells.^18^

In our work, we address these limitations, and show how the new framework we propose enables an improved quantitative perspective to the physiology of viral infections, paving the way for new *in silico* exploration of drug targets and multi-scale mechanistic models. Specifically, by adding a viral dynamics module to our framework for computation of microbial ecosystems in time and space (COMETS) (**Fig. 2**), we enable spatial (and potentially spatio-temporal) simulations of viral infections. At the same time, we redefine the objectives for the host-virus system in a way that is both mathematically tractable and biologically significant. The core idea is that a single ATP partitioning parameter controls at any given time the degree to which the virus has the capacity to hijack the host’s metabolism towards its goal. By handling the ATP allocation dynamically our framework can more accurately represent competition, and simulate infections over time to account for available nutrients and cell maintenance/death (**Fig. 2**). Our framework can thus answer questions regarding the role of resource allocation, specifically ATP allocation, on the death dynamics of the host cell and virus biomass production, which reveals the interplay between cell death and virus proliferation. In addition to parametrizing our model using realistic physiological parameters, we validated our modeling strategy with an improved SARS-CoV-2 infected iAB-AMØ-1410 macrophage model from Renz et al.^1,19–21^ Importantly, by incorporating the lipid component (in addition to the proteins) in the viral biomass, our version of the model improves the predictive capacity and suggests new possible drug targets. We envisage that COMETS can grow as a broadly usable platform for modeling any host/virus system, with adjustable parameters that can provide valuable insights on the role of metabolism in the dynamics of viral infections.

**Figure 2:**
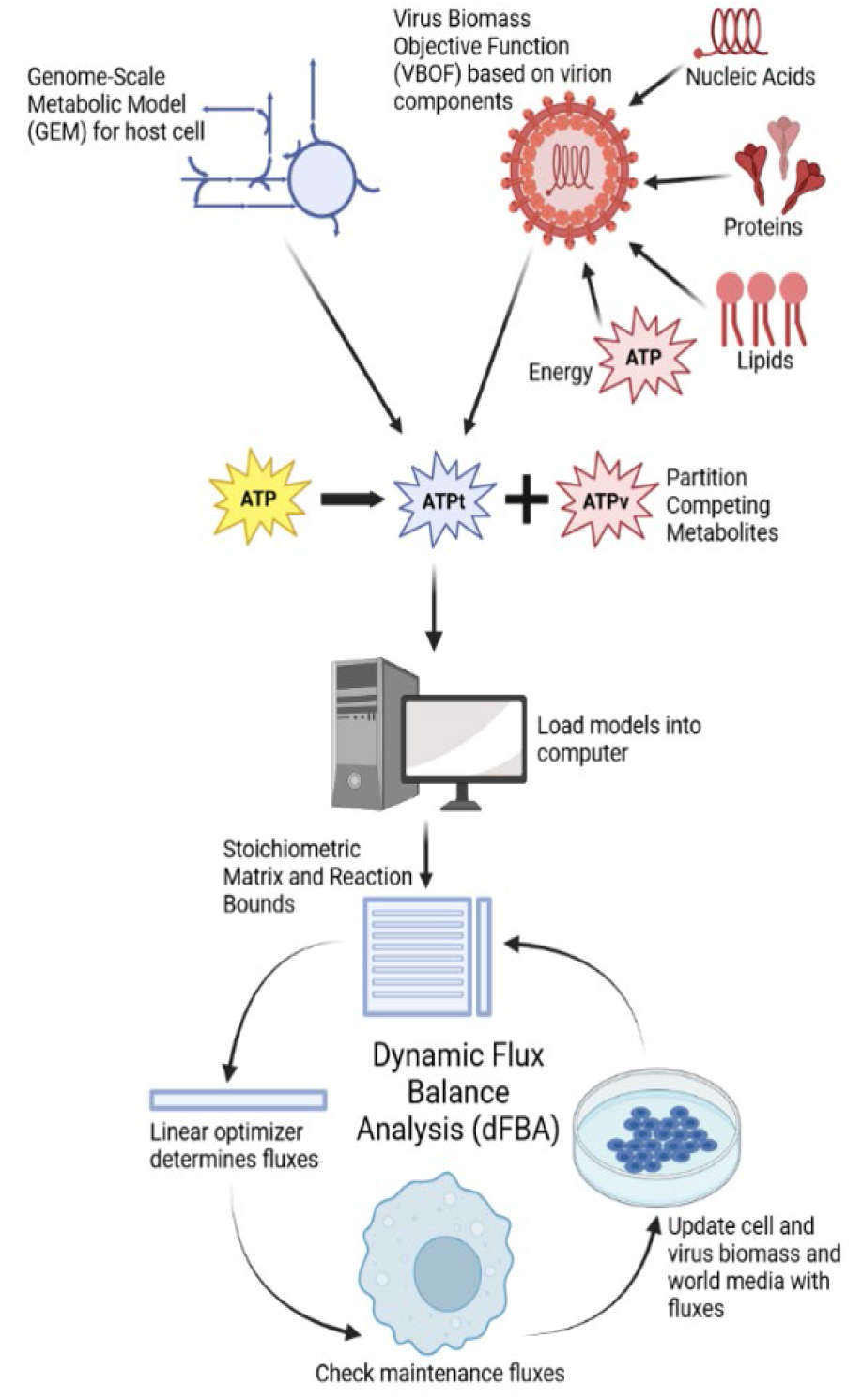
Framework of COMETS. It combines a host cell genome-scale model with the VBOF reaction representing virion synthesis from nucleic acids, proteins, lipids, along with the energy cost to combine the elements. COMETS then partitions competing metabolites and loads the models to run dFBA. Repeated iterations of FBA allow COMETS to simulate the cell infection over time, with the maintenance flux feature determining how many cells survive.

## Results

### Implementation of host-virus ATP competition in COMETS

Our viral module for COMETS simulates the metabolism of a homogeneously infected tissue of human cells through the use of genome-scale metabolic models (GEMs) (**Fig. 2**). Upon infection, the virus hijacks the host and reappropriates metabolites for its own reproduction.^22^ At the core of the competition between virus and host is hierarchical tree of metabolic processes that are encoded in the optimization step of COMETS for host-virus systems (**Fig. 3**). Since the host cell is typically not growing, but maintaining its homeostatic state through ATP-costly processes, we set the host objective to be the ATP maintenance reaction. Usually, in human FBA models, this reaction is assumed to have a fixed value, corresponding to the amount of ATP spent for any non-growth associated process that is not part of the metabolic stoichiometry itself (e.g. gradient maintenance, protein turnover, motility, etc.). Here, crucially, we assume instead that the human cells will try to maximize this flux (up to a certain upper bound) in order to stay alive. The portion of maintenance flux that can be satisfied translates into the portion of the human cells that can survive, while the remaining biomass (unable to satisfy maintenance) is bound to die. Thus, non-proliferating metabolism is modeled by satisfying maintenance reactions necessary for survival, particularly the energy cost associated with ATP hydrolysis.^23^ Meanwhile, the virus’ objective remains the production of virions, which is modeled by the maximization of the viral biomass objective function. However, if the virus demands too much energy from the host, the simulated cell will eventually die, rendering the virus unable to further replicate as well. Addressing this competition mathematically requires a set of computation steps that extends existing stoichiometric strategies, using a simple, but efficient multi-step algorithm (see Methods). In the end, the solution to this resource allocation problem is still obtained by using linear optimization to calculate the fluxes for each reaction and updating the environmental molecular composition accordingly (see Methods).

**Figure 3:**
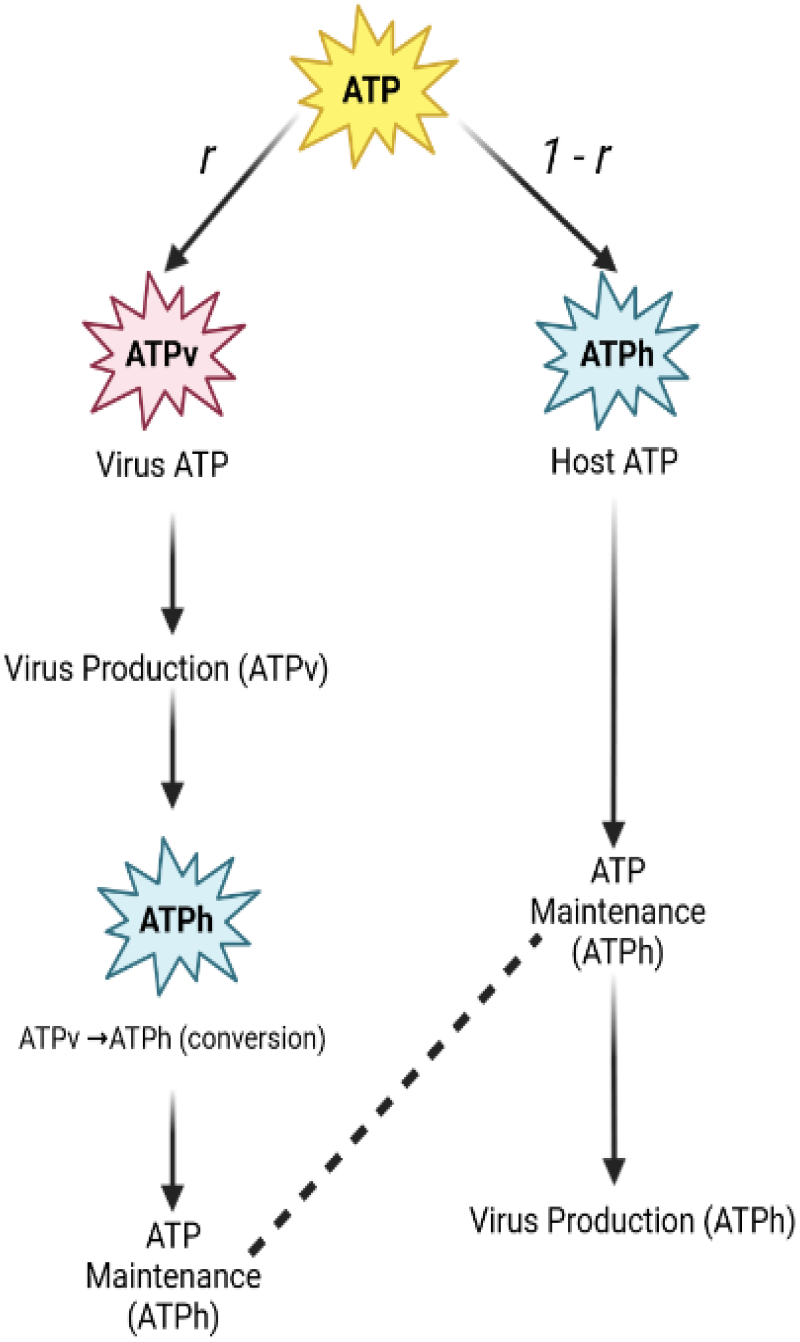
ATP is partitioned between the virus and the host objectives, creating the hierarchy shown. ATP is first divided into ATPv (ATP virus) and ATPh (ATP host) proportional to the partition ratio parameter *r*. ATPv is first consumed for virion production, and is then converted into ATPh for use in the maintenance reaction. Meanwhile, ATPh first goes towards host maintenance, then to the secondary Virus Biomass Objective Function (VBOFb). The total virus flux is computed as VBOF + VBOFb.

### Lipid Transport is a Potential Target for Drug Treatments of SARS-CoV-2

The SARS-CoV-2 host/virus model utilized in this study is a modified version of the one proposed by Renz et al.^1^ In this work, a SARS-CoV-2 specific Virus Biomass Objective Function (VBOF) was added to a human macrophage GEM. The virus biomass formulation in this model included the concentrations of amino acids, nucleotides, and energy required to synthesize a virion, but lacked calculations for the proportion of lipids. Using virus data from Das et al.’s simulations,^6^ we incorporated lipids into the computation of the virus biomass (see Methods section).

As a first step in our calculations, we performed standard FBA to determine the flux and growth rate impact of adding lipids in the virus’ biomass. We found that this addition of lipids had a nontrivial quantifiable effect on FBA calculations; most notably, the virion biomass production rate per cell (VBOF flux) increased by 29.6% compared to VBOFs that lacked a lipids component, and a total of 227 reactions were altered. The sum of absolute values of all host fluxes increased by 32.7% compared to VBOFs that lacked a lipids component, with their individual magnitude differences plotted in Figure 4. Most reactions kept the same flux directionality as in the previous viral model.

**Figure 4:**
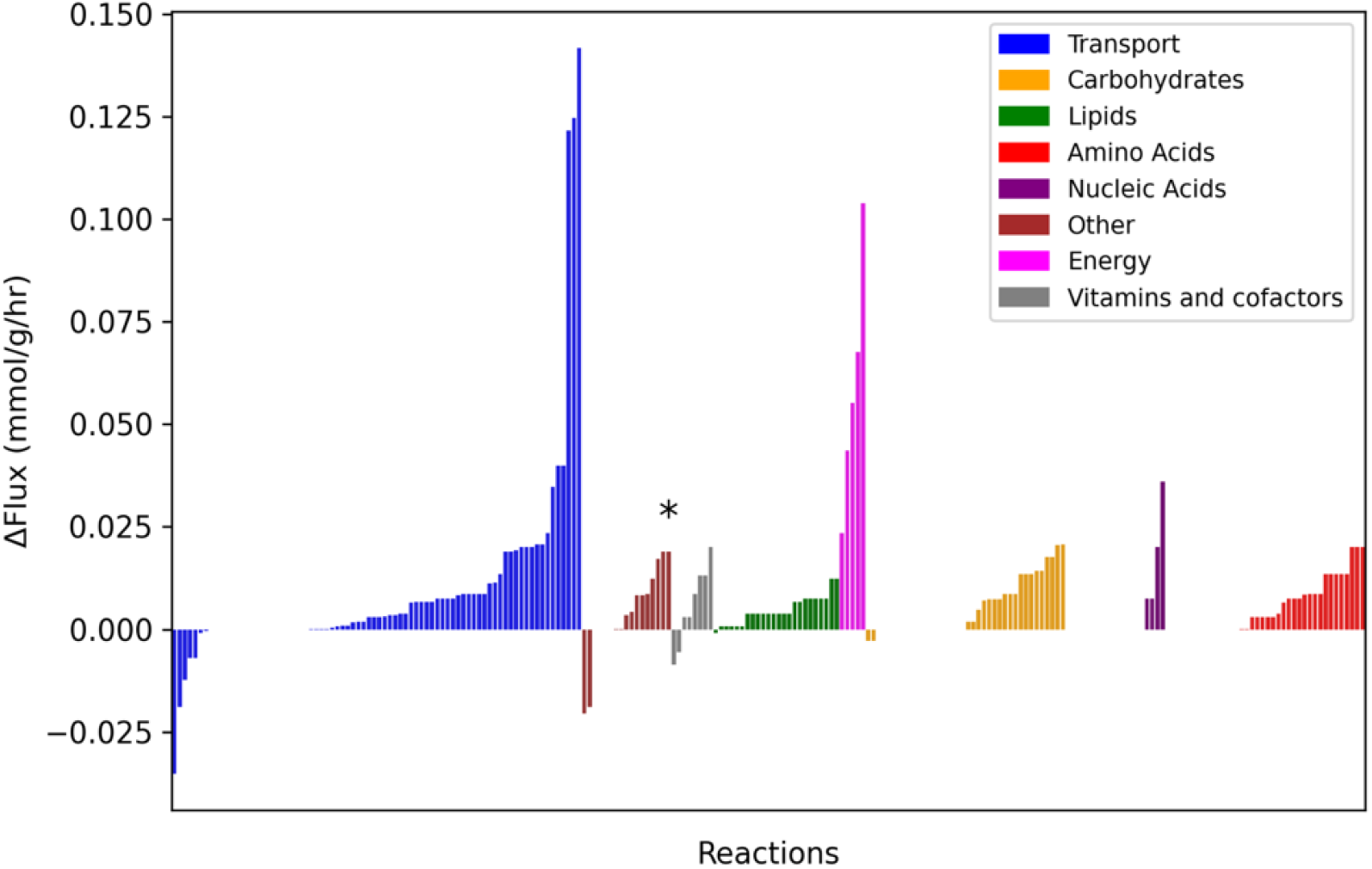
The inclusion of lipids in virus biomass computation results in a significant activation of specific host metabolic reactions. The bar graph depicts the nonzero differences in flux magnitude after including lipids. Reactions were colored by the pathways they belong to and the starred reaction, methylenetetrahydrofolate dehydrogenase, indicates the only reaction to reverse.

The effect of adding lipids on the viral biomass is also manifested in the different utilization of specific metabolic pathways. In particular, upon grouping the most drastically modified fluxes (with/without inclusion of lipids in biomass) into eight categories (lipid metabolism, carbohydrate metabolism, amino acid metabolism, nucleic acid metabolism, vitamin and cofactor metabolism, transport, energy production/consumption, and other), we found that transport reactions were altered the most (on average 0.41% per reaction), followed by energy production/consumption (∼5.80%), and then amino acid (∼2.87%), carbohydrate (∼1.11%), and lipid metabolism (∼3.42%) (Fig. 5). Overall, we observed significant changes in multiple systems, justifying the importance of factoring lipids into virus biomass.

**Figure 5:**
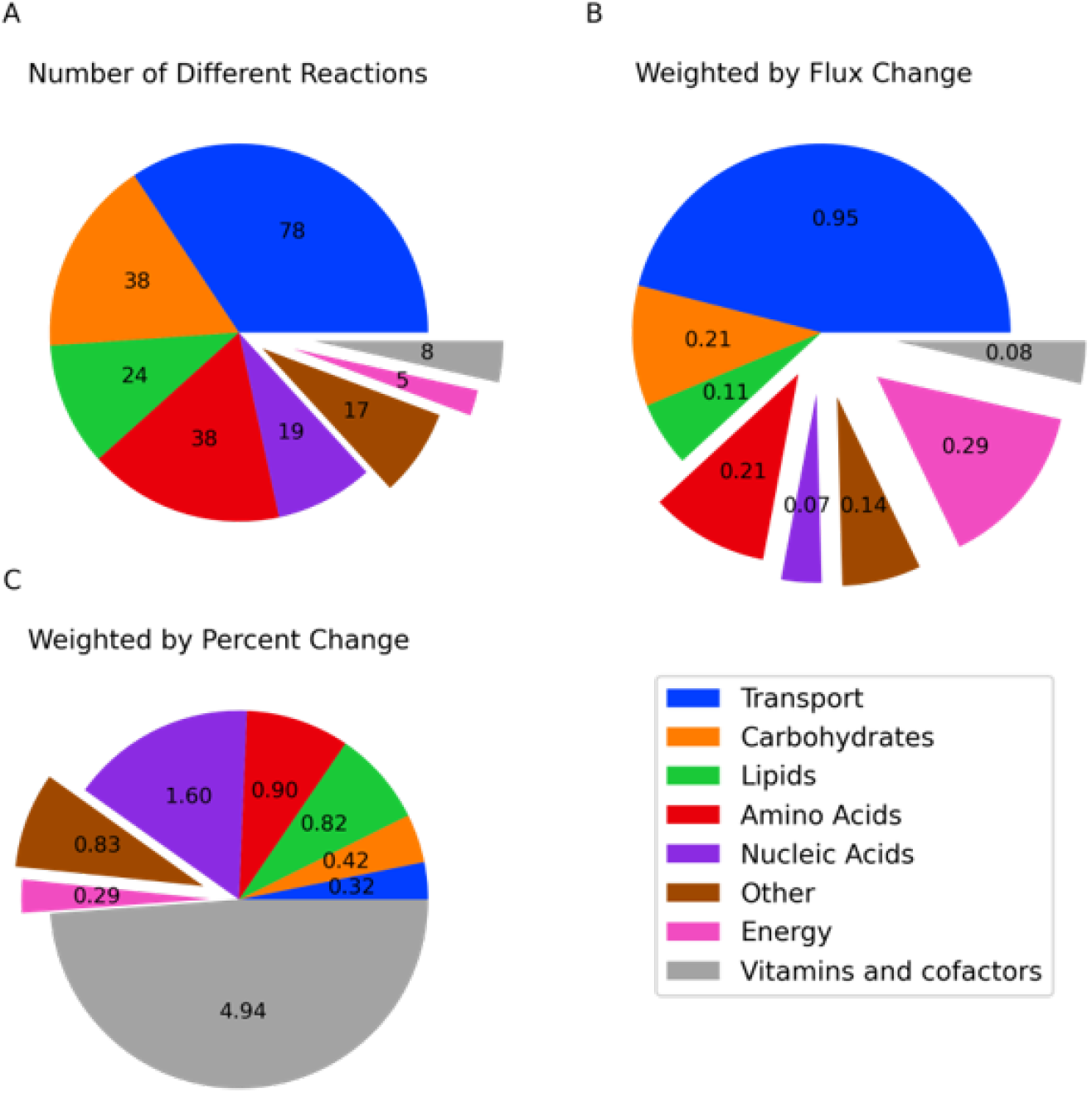
Inclusion of lipids into virus biomass computation results in significant alteration of host metabolic pathways, particularly among transport and energy related systems. Pie charts depict the differences between the lipids and no lipids models for different categories of reactions. A) is weighted by the number of reactions that changed B) by the amount the fluxes changed by, and C) by the percent change in the fluxes.

The implementation of a more accurate viral biomass composition allowed us to revisit this approach’s application to predicting possible drug targets, based on their modulation of human enzymes beneficial to the virus.. As done in prior work,^1,3^ we used FBA to simulate the effects of possible drugs by systematically blocking (i.e. setting to zero) the flux of each reaction in the human model, mimicking either the binding of the drug to the corresponding metabolic enzyme or the drug silencing transcription of the enzyme’s gene. To find reactions whose inhibition would point to relevant COVID-19 drug targets, we determined all reactions in our macrophage GEM that, when knocked out, would inhibit the virus’ growth without affecting host biomass pathways.

Our simulations identified five potential reactions to target for drug treatments of COVID-19: Guanylate Kinase, Sphingomyelin intracellular transport, Phosphatidylcholine scramblase, Methionine synthase, and 5,10 methylenetetrahydrofolate reductase. Interestingly, the Guanylate Kinase finding confirms a previous result from Renz et al.^1^ while sphingomyelin intracellular transport and phosphatidylcholine scramblase are novel targets that resulted from our inclusion of lipids in the virus membrane. In our simulations, the knockout of any of these three reactions completely halted the production of virions. Meanwhile, the other two reactions, belonging to amino acid pathways, were only able to halve the virus growth rate and were not dependent on our addition of lipids into the model. The viability of the sphingomyelin and phosphatidylcholine targets is assessed in more detail in the Discussion.

Ultimately, all of these results indicate that our addition of lipids in the virus biomass significantly affects the fluxes required to synthesize virions and the growth rate for the virus biomass.

### ATP Allocation Has Profound Effect on Host-Virus Dynamics

We next sought to use the human cell model and improved viral biomass composition to address the time-dependent competition for resources between host and virus. In particular, we took advantage of dynamic FBA (dFBA, through the COMETS framework)^24^ to simulate the competition for ATP between non-proliferating human cells (whose ATP is mainly used for maintenance and other non-growth associated metabolic functions)^25^ and viral particles geared towards maximizing their rate of reproduction at the expense of the host. In order to systematically explore the outcome of this tradeoff, we introduced in our model a parameter (the partition ratio, *r*) which controls the proportion of resources allocated to the virus relative to those allocated to the host. Thus, the fraction *r* of ATP will be prioritized for the virus, with excess given to the host while the remaining (*1 - r*) fraction of ATP will first go to the host with any leftovers available to the virus (see Fig. 3).

Our dFBA simulations were set to recapitulate growth of viruses in a population of non-proliferating human cell hosts, under a stable buffered amount of nutrients (see Methods). Parameters used in the simulations were either inferred from experimental studies or set to mimic *in vivo* conditions of human blood.^1,5^ Upon running simulations systematically for a fixed host ATP maintenance requirement and different degrees of viral “hijacking of ATP” (i.e. different values of *r* between 0 and 1), we obtained predicted values for growth/death rates and total biomass of both the host and the virus over time. As shown in Figure 6 when the virus utilizes only 10% of the ATP (*r* = 0.1) all the human cells survive the entire simulation. Furthermore, the constant virus growth flux and constant host biomass results in the virus biomass growing linearly with no bounds. When the virus reappropriates 30% of ATP (*r* = 0.3), viral growth flux increases, but it also causes the host cells to die exponentially since the ATP maintenance flux is forced below the required value. These factors result in the virus biomass reaching an upper limit because there are less cells to produce the virus as they die off. The simulation ends after around 2.5 hours, once all human cells are dead. The final case, where the virus utilizes 50% of the ATP (*r* = 0.5), is similar to the case when *r* = 0.3. However, the human cells die even faster (surviving only a little over 1 hour) and produce less total virus biomass despite a marginal increase in the virus growth flux. These results showcase the value of combining dFBA and a tunable ATP allocation, towards creating human infection models that can capture the metabolic tradeoff faced by the virus and its host.

**Figure 6:**
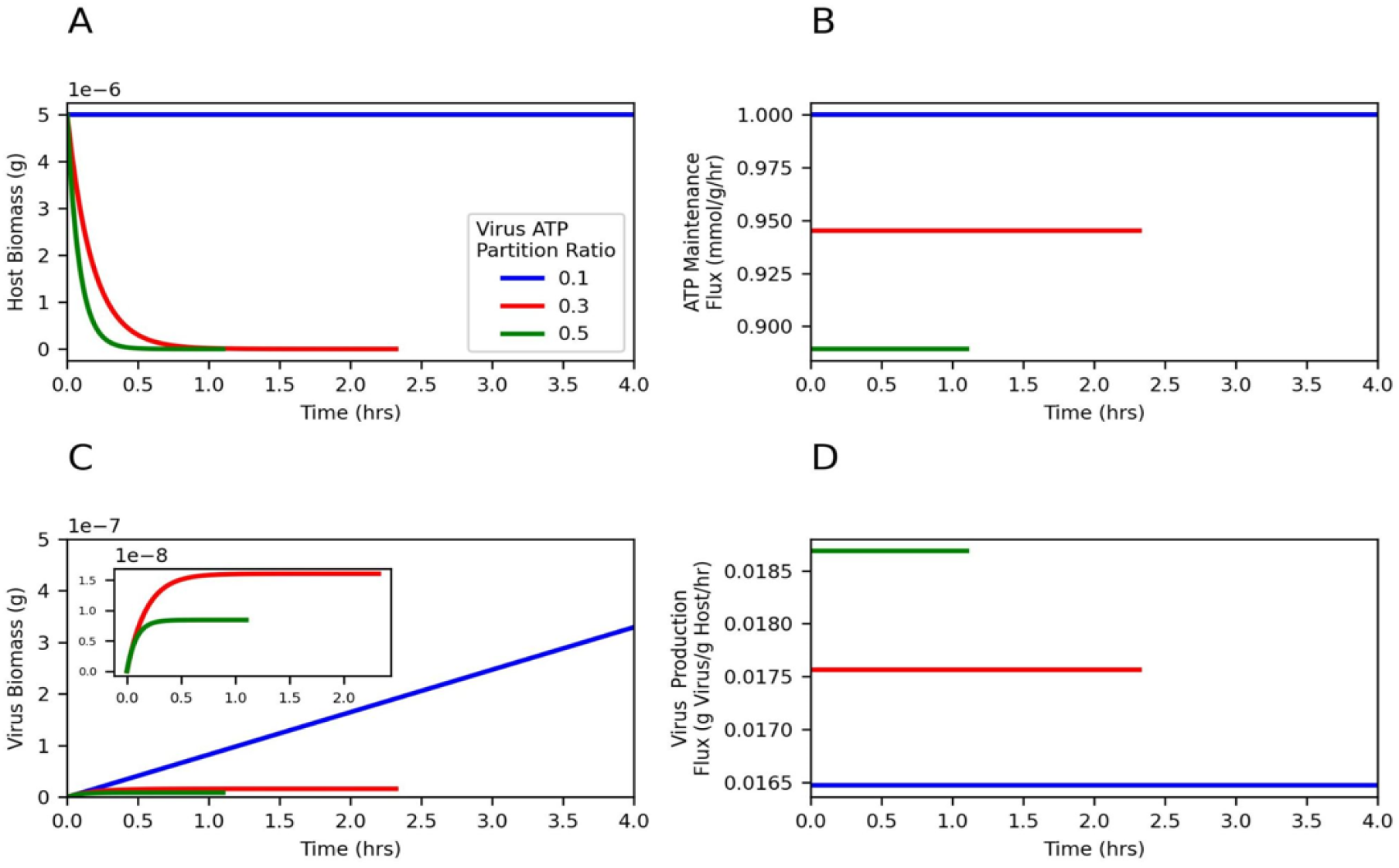
COMETS dynamic simulations show that the partition ratio has a profound effect on host-virus dynamics, where a higher proportion of ATP allocated to the virus (ATP Partition Ratio) leads to faster host death (A) and reduced virus biomass (C), despite the marginal increase in the virus production flux (D). The host death rate is proportional to amount of ATP requirement not satisfied which means a lower flux results in a faster death (A, B) while the virus grows proportionally to both host biomass and the virus flux. Subplots comparing three simulations with different partition ratios (*r* = 0.1 [blue], 0.3 [red], 0.5 [green]) and ATP requirement of 1 mmol of ATP/g/hr. The plots stop when the host biomass decreases to zero.

We then used our framework to more systematically investigate the result of all possible partition ratio values *r* on the final virus biomass yield as well as other factors that would contribute to the virus growth. Figure 7 graphs these effects, showing the host lifespan, virus flux, and virus biomass for all *r* ranging from 0 to 1. The host lifespan was unlimited for low partition values, but once *r* crossed a certain threshold, the host lifespan quickly decreased to a smaller constant value. The virus production flux behaved in the opposite manner, with lower partition values resulting in a smaller constant flux and larger partition values resulting in a larger constant flux. The model predicts a short transition period between these two states. The final virus biomass graph follows closely the host lifespan graph (Figure 7). In the region of low virus ATP partition the host cells never died and the virus biomass is therefore unbound. For higher partition ratios the host maintenance demands can no longer be met forever, and the host lifespan and the final virus biomass are finite.

**Figure 7:**
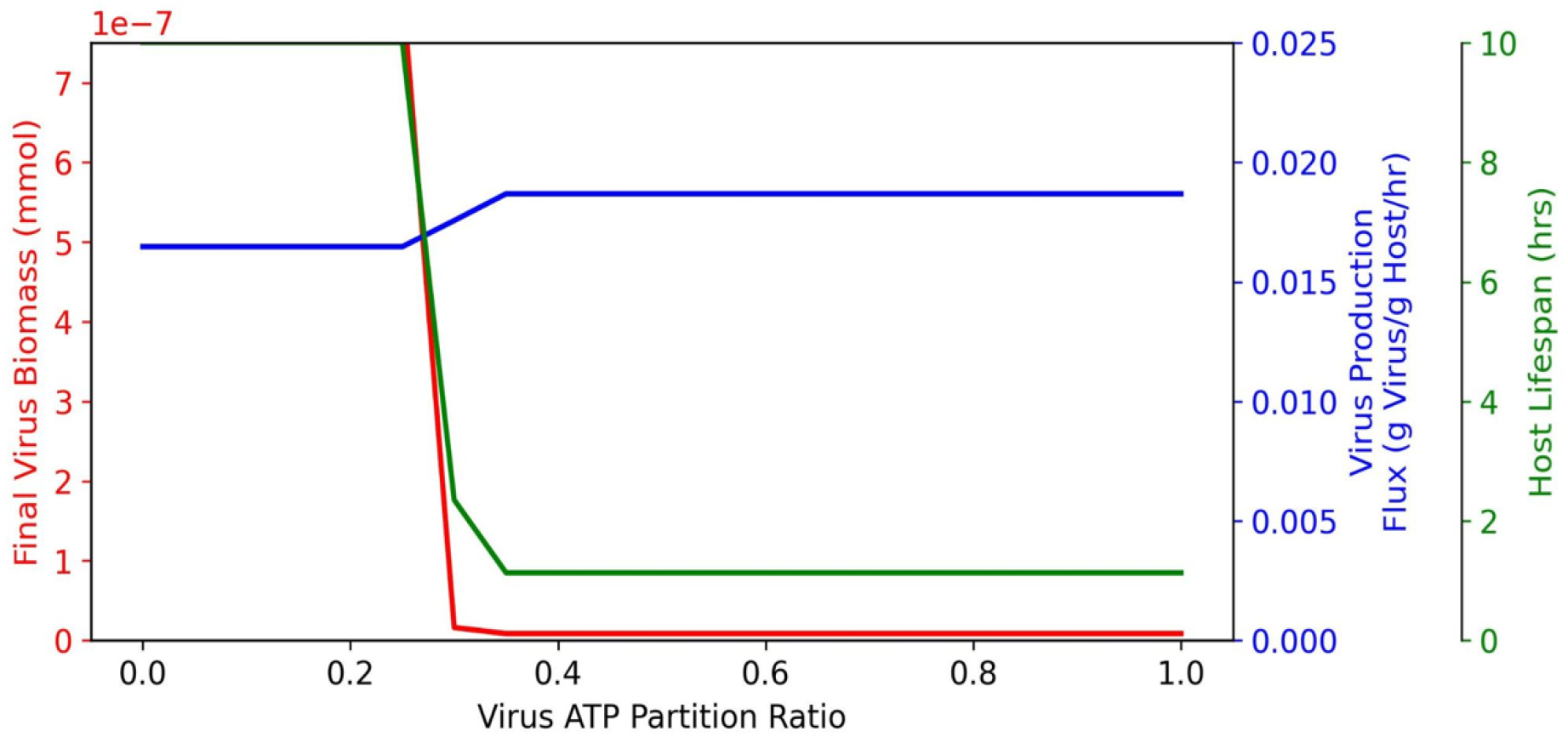
The final virus biomass is predominantly determined by the host lifespan, not the virus production flux. As more resources are partitioned to the virus (Virus ATP Partition Ratio), the host lifespan is reduced, with the host living transitioning from a non-limited/infinite lifespan to a finite one at ATP Partition Ratio threshold of 0.3 (red). Despite the virus production flux holding constant and increasing marginally at the same threshold (blue), the final virus biomass more significantly correlates with the host lifespan and similarly, infinite virus biomass is produced at a constant rate when the host lives forever.

Finally, we accessed the effects of varying the host ATP maintenance requirement (in mmol g^-1^hr^-1^) on the previously observed host/virus dynamics. As shown in Figure 8, there was a negative correlation between the ATP requirement and the partition threshold at which the cell lifespan begins to decrease; thus, higher ATP maintenance requirements increased sensitivity to the ATP partition ratio, resulting in the cell having a shorter lifespans and decreased virus biomass production at smaller partition ratios.

**Figure 8:**
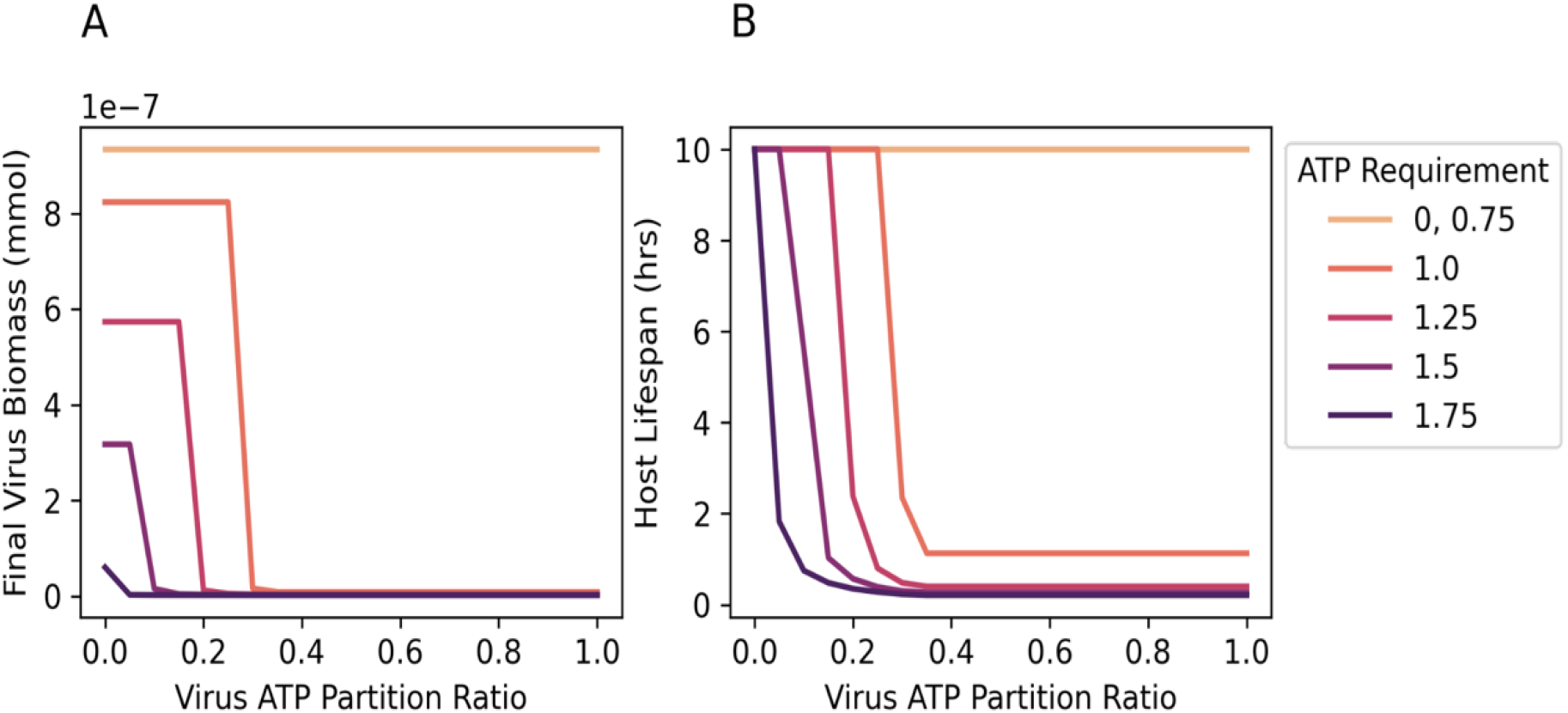
A higher host ATP requirement (indicated by darker colors) results in the host being more susceptible to lower partition ratios. The host lifespan was set to a maximum of 10 hours, and the final virus biomass was taken when at either when the host dies or at the end of the 10 hour simulation if it survived longer. As the ATP requirement increased, the threshold at which the cell survives over 10 hours decreased, and the final virus biomass was also strongly correlated with the host lifespan for all ATP requirements.

## Discussion

By enriching the viral objective function with experimentally-derived lipid content, recasting the optimization algorithm for host-virus interaction based on ATP competition, and embedding the virus-host stoichiometric system into a dFBA formulation, we have proposed a new avenue for performing simulations of viral dynamics in human cells.

The inclusion of the lipid component of the virus enables new testable predictions about possible drug targets. A potentially counterintuitive consequence of the incorporation of this additional resource in the viral biomass is the increase in viral biomass production rate, equivalent to a higher count of virions created per unit time (the VBOF coefficients are in millimoles of metabolite per gram of virus). The prior omission of lipids meant that a higher proportion of the virus was assumed to be composed of proteins and RNA. Thus, the cell actually needs slightly less amino acids, nucleotides, and ATP to synthesize a gram of virus than the previous model predicted, even though it had to now synthesize lipids as well. Because there is less contention in resources when synthesizing different metabolites, we observed a net increase in virus biomass production. This demonstrates the importance of including lipids in the virus biomass calculations.

COMETS gives researchers insight into possible metabolic changes associated with a viral infection; it provides a way to look for potential drug targets and can generate quantitative predictions of metabolic fluxes that can be compared to corresponding experimental measurements. Previous approaches^1,2^ to modeling viral growth using FBA have generally focused on a linear combination of fluxes (human + viral) as the objective, which leads to either the virus or the human cell “winning” all of the resources, with no middle ground where human tissue may struggle but survive, while the virus simultaneously replicates at a lower rate. This binary outcome is a direct outcome of the FBA method, which will always choose the state that gives the maximal numerical value for the objective. Here we specifically address the resource distribution between the host and the virus, by formulating an objective that can dynamically provide resources to both.

The dynamic simulations in Figures 6, 7, and 8 reveal interesting insights into viral replication strategies. Once the partition ratio is higher than a particular threshold the cell is drained of ATP for maintenance, and will die faster, limiting, in turn, the total virus biomass; the loss of cells that produce virions outweighs the increase in each cell’s ability to produce more virus biomass by devoting more resources to it. As seen in Figure 8, the graph of total virus biomass corresponds more directly with the host lifespan than the virus production rate.

Furthermore, the more ATP is required for a cell’s maintenance, the more competition there is and the more prone the cell is to dying: for example, if a virus partitions 20% of ATP for its own replication, a host that requires more than 1.25 mmol/(g⋅ hr) will die but a host with a lower maintenance cost will survive. Therefore, an increased host cell’s ATP demand (which may depend on cell type and environmental conditions), makes that cell more vulnerable to infections. Future work, along the lines of recent measurements of ATP competition during viral infection,^26^ could help test the model predictions, and fine tune parameters for the development of increasingly accurate simulations.

We should note that our model does not account for cell death from a variety of other reasons, such as when viruses burst out of cells during lysis, but focuses only on cell survival and death from a metabolic standpoint. The observed threshold which results in a sharp decrease in virus biomass (Figures 7 and 8) suggests that the optimal allocation strategy, from the perspective of the virus, would correspond to a partition ratio slightly under that threshold.

Our improved SARS-CoV-2 biomass composition allowed us to revisit the spectrum of possible drug targets. The model by Renz et al. identified guanylate kinase (GK1) as a potentially viable drug target to treat SARS-CoV-2^1^, and other works have shown the drug P1-(5’-adenosyl) P5-(5’-guanosyl) pentaphosphate (Ap5G) as a candidate for inhibiting GK1.^27,28^ In addition to recapitulating this result, we identify two new targets that have the potential to prevent or stunt the viral growth.

A first possible target we identified is sphingomyelin intracellular transport, which has been found to be an important pathway in influenza infections,^29^ another respiratory virus. In order for the influenza virions to bud off the vesicle, they rely on a variety of factors for intracellular vesicle transport such as the concentrations of lipid rafts composed of sphingomyelin, cholesterol, and glycosphingolipids.^29^ Higher concentrations of sphingomyelin was correlated with enhanced viral infection, whereas lower concentrations inhibited it.^30^

Imbalance in sphingomyelin concentrations has also been shown to correlate with cancer, so many works have attempted to find inhibitors for key enzymes that transport sphingomyelin within the cell ^31^. One such protein is the Ceramide Transport Protein (CERT), which is responsible for trafficking the sphingomyelin precursor ceramide from the endoplasmic reticulum to the Golgi. CERT functions independently of vesicle transport but relies on ATP to provide the necessary energy. The HPA-12 ceramide-related inhibitor and its derivatives have been the most prominent inhibitors of CERT. However, through more recent visual screening approaches, researchers have also identified HPCB-5 as the first non-ceramide-related CERT inhibitor with similar potency to HPA-12. These drugs could potentially be adapted to treat COVID-19 as well, by decreasing the supply of sphingomyelin in the Golgi for formation of the virion membrane.^31,32^

Another potential lipid-related target reaction is phosphatidylcholine scramblase. Although little work on COVID-19 has focused on phosphatidylcholine, Zhang et al.^33^ found that the Brome Mosaic Virus, also a positive-sense RNA virus, stimulated phosphatidylcholine synthesis in the host metabolism. However, scramblases, particularly TMEM16F, have been shown to play a key role in SARS-CoV-2 pathogenicity. TMEM16F externalizes phosphatidylserine on the outer leaflet of the membrane which promotes cell fusion events, including spike-driven viral entry.^34^ Since the antihelminthic drug niclosamide (among many others) is known to inhibit TMEM16 family proteins, Braga et al. have proposed to use it as a potential COVID-19 treatment. Given that scramblases are nonspecific,^35^ it could be possible for TMEM16F to also play a role in internalizing phosphatidylcholine needed for viral replication. TMEM16F would then be a very promising target for drugs as it could reduce both virus pathogenicity and viral replication.

## Conclusion

By using COMETS, we implement dFBA simulation of viral infections over time based on a new framework for partitioning limited resources for competition and allocation. We applied our framework to simulate SARS-Cov-2 infected macrophages, explicitly including viral lipid concentrations for use in FBA calculations. We demonstrate that this inclusion significantly alters the fluxes and allows for the identification of two new possible drug targets. Finally we explore the viability of these targets, and demonstrate the effects of the host ATP requirement and partition ratio on virus biomass and host lifespan. Our work provides an improved mechanistic modeling tool for exploring viral strategies during infections and for suggesting potential therapeutic treatments.

One should keep in mind the limitations of the proposed approach. First, being based on FBA, our method is inherently restricted to the prediction of metabolic fluxes, and cannot provide insight into processes dependent on intracellular metabolite concentrations, including transcriptional and post-translational regulatory effects. Moreover, our simulations do not currently include details of the immune response, which plays a major role in fighting viral infections. Modeling these would require more complex models that record various intra- and inter-cellular processes, as well as different cell-types and substantial molecular details.

Second, our current model focuses on competition for one specific resource (ATP) but future versions could include competition for other resources needed for cell maintenance, repair, and metabolite turnover, including additional currency metabolites. In addition, our study is currently limited to a homogeneous (i.e. non-spatially structured) tissue composed of infected cells. However, given the capacity of COMETS to simulate spatially structured environments populated by different cell types and infection states, future iterations could extend our framework to explore interactions between host cells in advanced 3D models of tissues. We envisage that such extended models could help simulate the propagation of infections across different cell types in different layers potentially matching more realistic experimental scenarios and observations.

It is finally worth noting that while our work is entirely focused on the specific scenario of SARS-Cov-19 infection of human cells, the viral infection module we implemented in COMETS can be easily applied to study any viral infection, including the dynamics of phages in bacterial cells and microbial communities, with possible applications in the study of host-associated, marine or soil microbiomes.

## Methods

### Flux Balance Analysis (FBA) and Dynamic Flux Balance Analysis (dFBA)

The host-virus interaction is addressed here using dynamic flux balance analysis (dFBA). This approach, first introduced in 2002^36^ is based on iterating solutions of a Flux balance Analysis (FBA) problem at a series of time points. The standard formulation of FBA, described in detail before,^14,24,37^ relies on assumptions of steady state and optimality to compute a putative vector of metabolic fluxes for all reactions in the metabolic network of a given organism. The efficiency of FBA is due to the fact that it can be solved using linear programming.^11–14^

The notation used below is similar to that used in several prior FBA studies. We denote with *S* the stoichiometric matrix, whose element *S*_*i,j*_ encodes the stoichiometry of metabolite *i* in reaction *j*. The vector of fluxes *v* is required to satisfy steady state constraints, expressed by the equation *Sv* = 0. The objective function is expressed through a vector *c*, such that the flux combination to be maximized is *c*^⊤^*v*. The non-zero elements of the vector *c* can be interpreted as weights for each of the reactions contributing to the objective. In our viral model, the objective function is a combination of three reactions with significantly different weights, as described in detail below. Furthermore, each reaction is bounded by a pair of numbers (an upper *u* and a lower bound *l*), which encode thermodynamic irreversibility, nutrient availability, and other constraints.^24^ With this nomenclature, the FBA problem is described as follows:

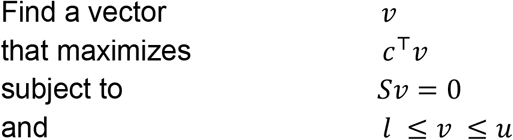

In order to implement simulations of time courses, in which intracellular metabolism is assumed to quickly approach steady state, we implement dynamic FBA (dFBA).^36^ dFBA stores information on the world media (i.e. the concentrations of different metabolites available to the cell). It then does repeated iterations of standard FBA with reactions bounded by the amount of nutrients available to compute the uptake fluxes and update the world media accordingly. The standard FBA supplies the growth rate of the biomass, as well as the uptake and secretion rates for the exchanged metabolites. The growth rate is an input for the dFBA algorithm that numerically solves the differential equation of the biomass growth over time. The extracellular metabolites in the media also can change dynamically with the rates of the exchange reactions.^36,37^ The upper bounds of the uptake rates for an extracellular nutrient are typically calculated with the Michaelis-Menten formula:

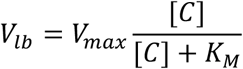

Given the biomass *B* and its growth rate *v*_*gr*_, and the uptake/secretion rates {*um*} of extracellular metabolites {*Q*_*m*_}, the dFBA algorithm solves the differential equations of the growth and change of extracellular metabolites concentrations:

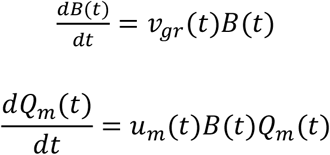

In this work we used the standard convention of *um* > 0 for secretion and *um* < 0 for uptake.

In this work, we modify the standard dFBA approach to mimic the conditions of a virus-infected macrophage in the human body. We assume that in the body the human cells are constantly supplied with nutrients, therefore setting conditions of constant concentrations of the extracellular metabolites. The dFBA algorithm is therefore modified in our model such that we keep the concentrations of the extracellular nutrients constant, i.e.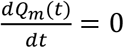. As a consequence, the upper bounds of the uptake rates also do not change over time (*um*(*t*) = c*onst*). We set the upper bounds of nutrient uptake fluxes according to the values proposed in Renz et al., which mimic the nutrient composition of human blood.

Our platform COMETS is an implementation of the dFBA methodology, written in Java, and accessible through a Python interface: cometspy. COMETS and cometspy are open source software, publicly available at https://github.com/segrelab/comets and https://github.com/segrelab/cometspy. They can be downloaded and installed from https://www.runcomets.org/.

### Human macrophage metabolic reconstruction

To simulate the host metabolism, we used the macrophage cell-specific model iAB-AMØ-141039 provided in reference Bordbar et al. (2010) with Renz et al. (2020) adding the SARS-CoV-2 virus objective. The underlying model of the general human metabolism is based on the Recon 1^15,38^ model, with a total of 3394 reactions and 2572 metabolites.

The model was grown on a medium with nutrients based on metabolites present in human blood.^39^ The media in the model is defined by setting the metabolite specific upper bounds of the uptake rates, specific to the type of cell (macrophage) that is modeled.^1^ Most importantly, the maximum uptake rate for glucose, typically the limiting nutrient, was set to 0.2718 mmol/(gr⋅ h).

### Modeling Viral Infection

#### Virus Biomass Objective Calculations

Since viruses lack metabolic pathways of their own and must utilize the machinery of the cell to replicate, they can be modeled as a virion production reaction integrated into a host GEM. Aller et al. define this reaction as the Virus Biomass Objective Function (VBOF).^5^ It uses proteins, nucleotides, and energy to produce viral biomass with coefficients for each metabolite, *Z*_*X*_, expressed as millimoles of metabolite per gram of virus (mmol/g). Let us denote by *X*^*TOT*^ the moles of *X* per mole of viral particles and by *M*_*v*_ the molar mass of the virus (the weight in grams of an Avogadro number of viral particles) where *M*_*v*_ is given by the equation:

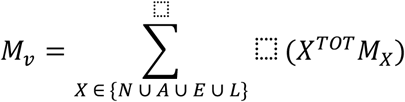

with *N* denoting the set of all nucleotides, *A* denoting the set of all amino acids, *E* denoting the set of molecules required for energy consumption, *L* denoting the set of all lipids that constitute the virus biomass, and *M*_*X*_ as the molar mass of metabolite *X*. Then *Z*_*X*_ in mmoles per gram for each metabolite *X* can be computed by:

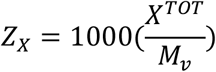

where *X*^*TOT*^ is the count of metabolite *X* in each virion and *M*_*v*_ is the molar mass of a virion.^1^ Note that Aller et al.^5^ as well as other FBA works on virus studies excluded lipid metabolites from the VBOF calculations due to a lack of data.^1–3^ Our work is, to the best of our knowledge, one of the first viral FBA models to incorporate accurate lipid synthesis.^4^

#### Inclusion of Lipids in Virus Biomass

We use data from the Das et al. study on determining SARS-CoV-2 membrane stability in the presence of ethanolic disinfectants to approximate the membrane composition.^6^ Since coronaviruses have been found to bud off the Endoplasmic Reticulum-Golgi Intermediate Compartment (ERGIC), they assumed the viral membranes would be composed of the same lipids as the ERGIC. From this, they estimated that the membranes include sphingomyelin and phosphatidylcholine in a 1:1 composition ratio.

The formula for Aller et al.’s VBOF coefficients requires the total count of each type of lipid, which we can determine based on the virion’s size. The average SARS-CoV-2 virion is estimated to have a diameter of 100 nm^40,41^ which means the surface area would be approximately 31,415 nm^2^. Das et al.’s studies revealed an average area of 56 Å^2^ per lipid.^6^ Since there are two leaflets in the viral membrane, there are around 112,200 lipids in total, 56,100 molecules of sphingomyelin and 56,100 molecules for phosphatidylcholine. From this, we estimated the coefficients for both lipids in the VBOF to 0.2023 mmol/g.

Comparing the inclusion and exclusion of lipids in virus biomass: We compared the two models using COBRA,^42^ the standard toolset for FBA simulations of genome-scale models. We directly modified their VBOF to include lipids cost. Both the adjusted and original model were run using parsimonious FBA with the maximizing the VBOF as the only objective. Parsimonious FBA, in addition to maximizing the objective, minimizes the sum of the absolute values of all fluxes. The fluxes for each reaction were then recorded from both simulations and compared.

#### Partitioning Resources for Virus Host Competition

We use the VBOF as the objective for the virus, while we use ATP maintenance for the host. Both objectives require cytosolic ATP, but the hierarchical nature of FBA means only one reaction will receive it. This means that if a sequence of optimizations is run, the first one on that sequence will obtain it all. This problem cannot be remedied with, for example, change of the weight coefficients in the objective function. Changing the linear combination weights will only switch which reaction receives the ATP, preventing the virus and host from coexisting. To solve this, we partition cytosolic ATP using a pseudoreaction and generate a new hierarchy shown in Figure 3. A certain fraction of the resource will always be explicitly allocated to each objective reaction. In other words, part of the ATP will be allocated to the host maintenance and part to the virus biomass reaction. If one of the objective reactions cannot uptake all of its partition, only then will its resources be redirected to the other objective. Thus, resources of one partition will always go to the virus first and only its leftovers will go to the host maintenance, while the resources of the second partition will always go to the host maintenance first and only its leftovers will go to the virus. Biologically, this is equivalent to assuming that any excess ATP not being used, will be consumed by whichever objective needs it.

We achieve this partitioning by manipulating the stoichiometric matrix by introducing two new metabolites—ATP_h (ATP Host Maintenance) and ATP_v (ATP Virus), and modifying our objective reactions so they are consumed by the ATP maintenance reaction (ATPm) and the VBOF respectively. Then we introduce 2 new pseudo-reactions:

1. VBOFb: a copy of the VBOF except instead of consuming ATP_v, it consumes ATP_h; this provides an alternative pathway of virus production once ATP maintenance of the host has been satisfied. This means that the virus can utilize excess ATP left from the host for its own growth.
2. ATP Conversion: ATP_v → ATP_h; enables excess ATP_v not used by the virus to go to host maintenance by converting it into ATP_h. The advantage of the conversion is that ATP maintenance (ATPm) can remain as a single reaction instead of the sum of two reactions (as the VBOF is forced to be).

The dFBA algorithm for simulating virus infection is a version of the standard dFBA modified in order to maximize two objectives simultaneously, virus growth and host maintenance (VBOF and ATP maintenance). Here below are the steps performed by our modified dFBA at every cycle to implement newly proposed resource partition:

1. Set the bounds of nutrient uptakes *V*_*l*_ for each extracellular metabolite based on its concentration using Michaelis-Menten equation as in standard COMETS dFBA algorithm.
2. Optimize (maximize) a linear combination of three fluxes (VBOF, ATPm, VBOFb) to capture the hierarchy of possible fates of ATP. In particular we define the host cell maintenance and virus biomass growth objective function as follows: *W*_*v*_ *VBOF* (*consumes ATPv*) + *W*_*h*_ *ATPm* (*consumes ATPh*) + 1 *VBOFb* (*consumes ATPh*).Here we use *W*_*v*_ and *W*_*h*_ to 10^6^ and 10^3^ respectively in order to impose a priority of objectives relative to the VBOFb which is weighted by 1. Thus, by the nature of the linear combination of objectives, the optimization algorithm will satisfy objectives in the following order: VBOF, ATPm, and VBOFb.
3. For the optimal values of the fluxes computed in the previous step, calculate the total virus growth rate as *v* = *VBOF* + *VBOFb* and estimate the total virus biomass accumulated at this rate over a time period Δ*t*, according to the formula Δ*m*_*v*_ = v ⋅ m_*h*_ ⋅ Δ*t*.
4. Check if the ATP maintenance requirement for the host cell is satisfied, i.e. assess the value of ATPm for the above computed solution. The requirement is satisfied if *ATPm* ≥ v_*min*_. If it is not satisfied, we assume that the human cells will die in proportion to the fraction of maintenance not satisfied. In practice, this means that we reduce the host biomass by an amount proportional to the ratio *v*_*m*_/*v*_*min*_:

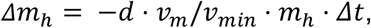

where *d* is a parameter that we set to *d* = 100 *h*^−1^. This parameter which sets the time scale for the death process was chosen conservatively so as to match experimental observations of infected hosts cell dying rate in the range of several hours.^43^

Our objective is hierarchically defined as follows (from highest to lowest to priority): VBOF, ATP maintenance, VBOFb, creating the structure shown in Figure 3. This scheme then enables us to model the competition of resources by allowing ATP to be consumed by different reactions in different orders. ATP_h will first be used in ATP maintenance because it cannot be uptaken by the VBOF. Once ATP maintenance reaches its maximum flux, the leftover ATP_h will be devoted to VBOFb, the next highest objective in the hierarchy. Meanwhile, ATP_v will first be used in the original VBOF because it has the highest priority. Once the VBOF is maxed out, ATP_v will be converted into ATP_h which in turn is used in ATP maintenance. The total virus flux is then calculated by the fluxes of VBOF + VBOFb.

#### dFBA within COMETS Simulates Maintenance Fluxes with Cell Death

This modified GEM provides the input for dFBA simulations,^24,36^ which predict the fluxes and extracellular metabolite abundances at each time step ; specifically, we use parsimonious dFBA, which assumes that, among all possible fluxes that maximize the objective function, the cells adopts those which are most parsimonious, i.e. also minimize the sum of reaction rates. In our revised implementation of dFBA, we also add a feature to account for cell death based on the extent to which the maintenance reaction flux is not supported. Traditional approaches encoded maintenance reactions with a strict lower bound, resulting in an infeasible solution if the maintenance flux cannot be satisfied.^44–46^ However, our model removes these constraints and a portion of the host biomass dies instead; the host only retains the percentage of biomass it can sustain. Thus, after every time interval 1/*d*, if the maintenance flux *v*_*m*_ is less than the minimum ATP requirement *v*_*min*_, only a fraction *v*_*m*_/*v*_*min*_ of the original host biomass will survive.

Once the host cell biomass reaches zero, we end the simulation since all of the cells have died. Furthermore, since we assume that a cell will not consume more ATP than required to survive, we constrain the upper bound for the maintenance reaction to *v*_*min*_.

The virus growth rate will depend on both the VBOF flux and the host biomass, as the virus replication is proportional to the number of living, infected cells. Thus, the change in virus biomass at each time step is given by:

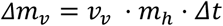

where *m*_*v*_and *m*_*h*_ are the masses of the virus and host cell aggregate respectively, *v*_*v*_ is the flux of the VBOF in mmol/g/hr, and Δ*t* is the time step change for each dynamic FBA iteration in hours.^43^

## Key Resources Table

| Resource | Source | Identifier |
| --- | --- | --- |
| COBRA | Ebrahim et al. | http://opencobra.github.io/ |
| COMETS | Dukovski et al. | https://github.com/segrelab/comets |
| Gurobi Optimizer | Gurobi Optimization, LLC. | https://www.gurobi.com/ |
| BiGG Database | King et al. | http://bigg.ucsd.edu/ |
| iAB-AMØ-1410 macrophage & SARS-CoV-2 model | Renz et al. | https://www.ebi.ac.uk/biomodels/MODEL2003020001 |

### Determining Potential Knockout Targets

Using the new model that accounted for lipids in the virus biomass, we ran parsimonious FBA using COBRA, optimizing only the VBOF to determine the maximum virus replication flux. Then, for every reaction in the host GEM, we restricted its upper and lower bound to 0 to model the effect of knocking out the reaction from the system, and ran parsimonious FBA again to determine the virus replication flux. If this flux was less than the maximum flux, we recorded it as a candidate.

Of those candidates, we ensured they did not affect any potential host biomass growth. The maximum host biomass flux was computed with another parsimonious FBA run but without any reaction knockouts and optimizing the host biomass instead. We then knocked out each candidate reaction and ensured that the host biomass did not deviate from the maximum growth rate. Thus, the five reactions we found restricted the virus flux to less than its maximum but did not affect the host growth rate at all.

## Acknowledgements

This work was partially supported by the National Institutes of Health (National Institute on Aging - award number UH2AG064704, and National Cancer Institute - grant number R21CA279630) and by the National Science Foundation Center for Chemical Currencies of a Microbial Planet (C-CoMP publication #047). Author contributions Conceptualization, A.L., I.D., D.S.; Methodology, A.L., I.D., D.S.; Software, A.L.; Formal Analysis, A.L.; Investigation A.L., I.D., D.S.; Data curation, A.L.; Writing - Original Draft, A.L.; Writing - Review and Editing, I.D., D.S.; Visualization, A.L.; Supervision, I.D., D.S.; Project Administration, I.D., D.S.; Funding Acquisition, I.D., D.S.

## Materials and correspondence

Requests for materials and correspondence should be addressed to I.D. and D.S..

